## Supplementary figures and images for "Humans strategically shift decision bias by flexibly adjusting sensory evidence accumulation"

### Fig 2 sup 1

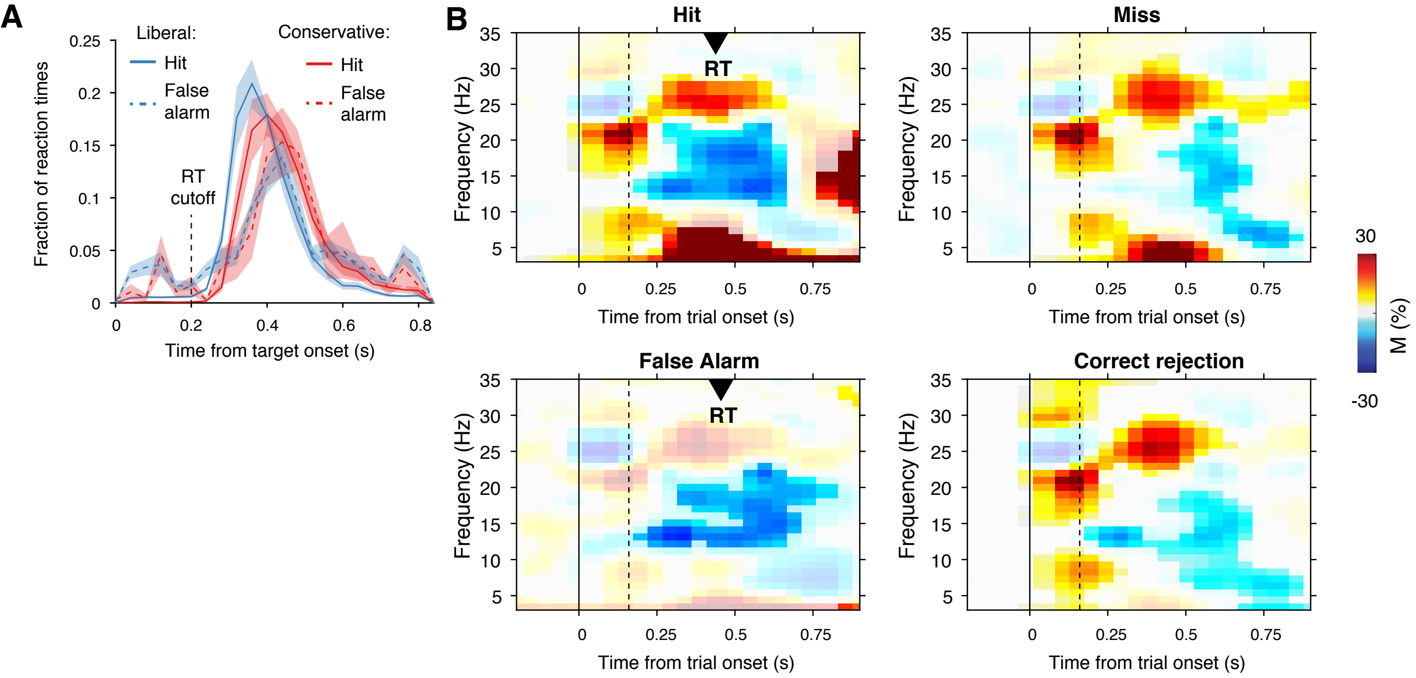

### Fig 2 sup 2

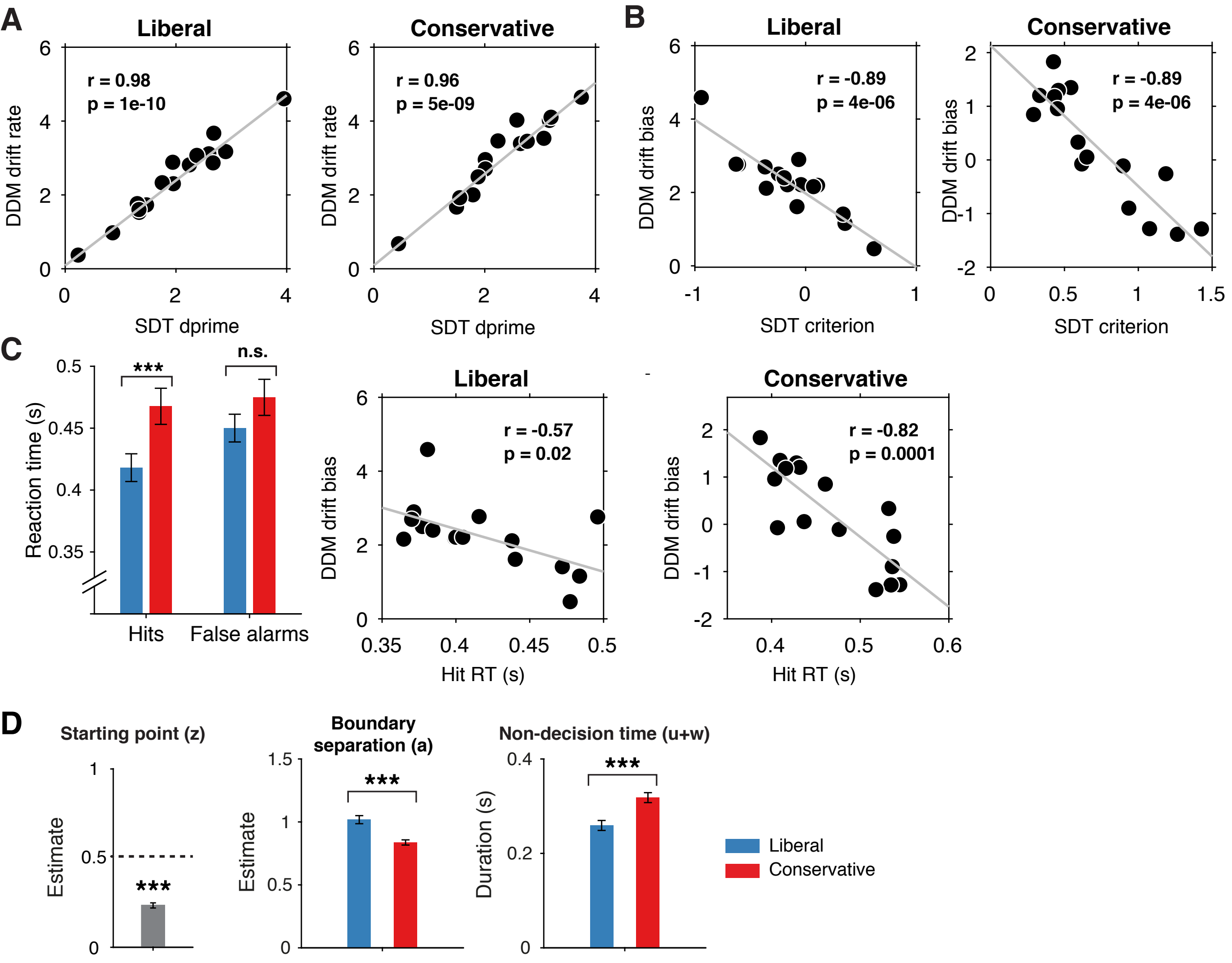

### Fig 2 sup 3

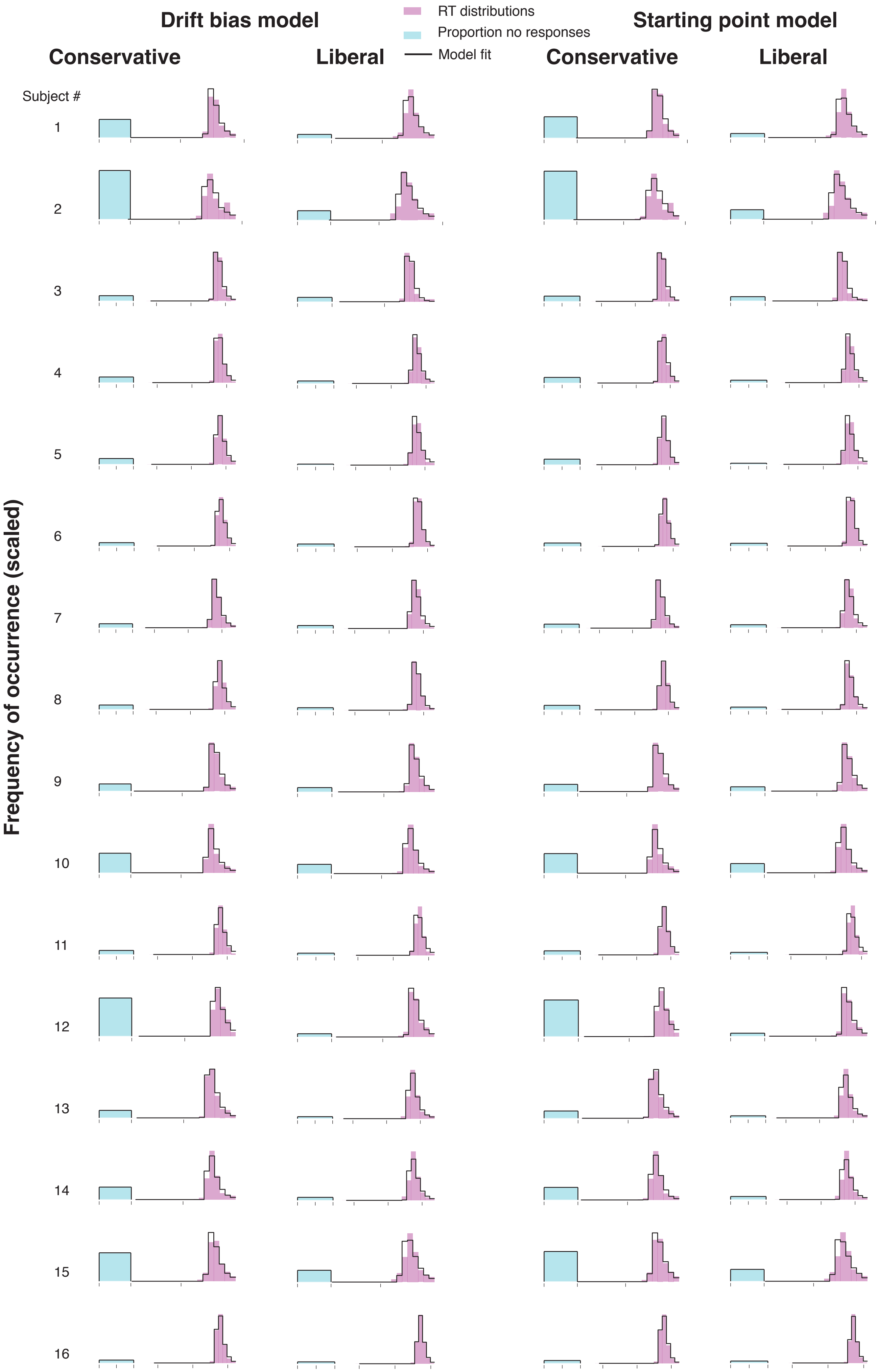

### Fig 6 sup 1

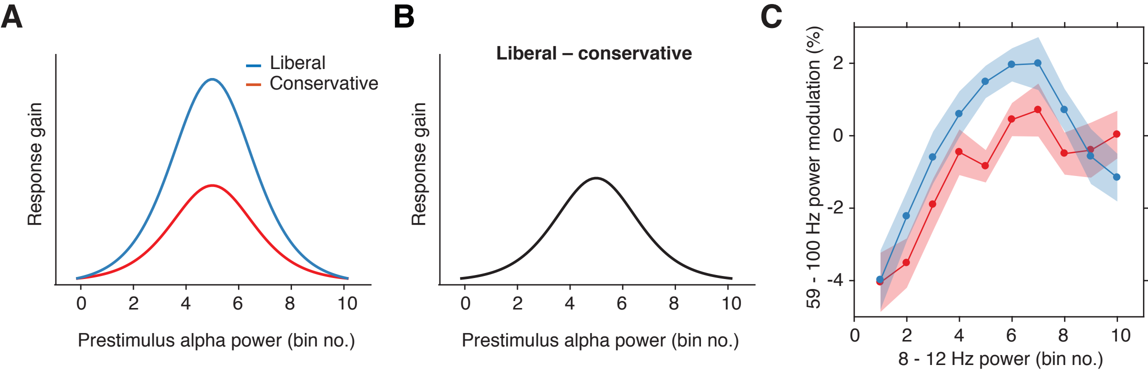

### Fig 7 sup 1

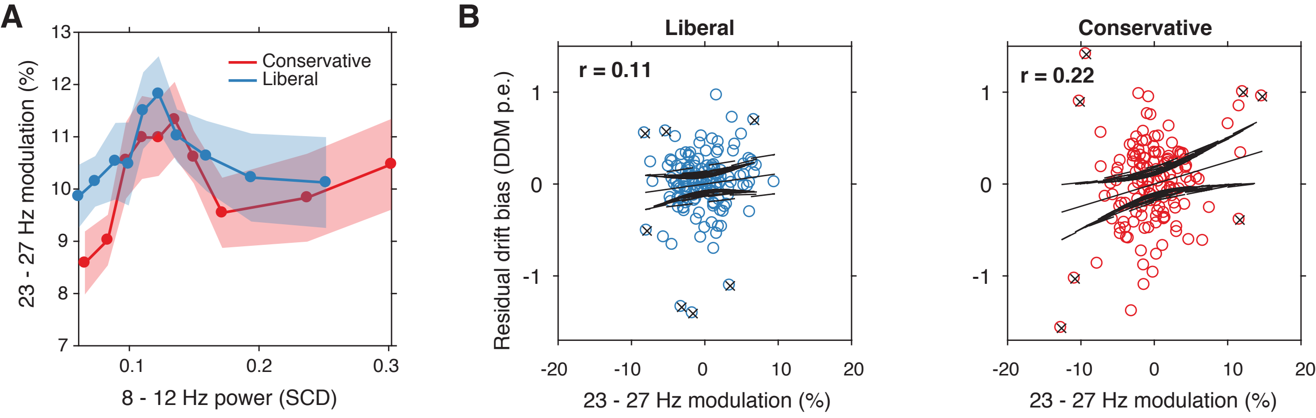
